# Cadmium-induced endocytosis of the broad spectrum root metal transporter of Arabidopsis

**DOI:** 10.1101/2022.02.21.481249

**Authors:** Julien Spielmann, Virginia Cointry, Julie Neveu, Grégory Vert

## Abstract

Iron is an essential micronutrient for plant growth and development. Under low iron conditions, Arabidopsis plants take up soil iron using the root iron transporter IRT1. In addition to iron, IRT1 also transports others divalent metals including cadmium that consequently accumulates into plant tissues and enters the food chain. *IRT1* expression was shown to be regulated at the transcriptional and post-translational levels by its essential metal substrates to maximize iron uptake while limiting the accumulation of zinc, manganese or cobalt. Here, we characterized the regulation of IRT1 by cadmium and uncovered a cadmium-mediated downregulation of IRT1 protein by endocytosis. A short term exposure to cadmium indeed decreased the celle surface levels of IRT1 through endocytosis and degradation. This is mediated through the direct binding of cadmium to histidine residues within the regulatory loop of IRT1. Moreover, we demonstrated that cadmium-induced IRT1 degradation uses ubiquitin-mediated endocytosis driven by the IDF1 E3 ligase. Altogether, this work sheds light on the mechanisms of cadmium-mediated downregulation of IRT1 and offers a unique opportunity to boost plant cadmium uptake in phytoremediation/phytoextraction strategies.

## Introduction

Iron (Fe) is an essential trace element for all living organisms due to its ability to easily loose or gain electrons. Fe serves as a major cofactor for a wide range of electron transfer reactions such as photosynthesis or respiration. Due to its propensity to precipitate in aerobic conditions or at alkaline pH under the form of insoluble ferric oxide/hydroxide, iron is not readily available in the rhizosphere and often limits plant growth. Fe deficiency reduces chlorophyll biosynthesis and photosynthetic activity, yielding leaf interveinal chlorosis and decrease in biomass (Briat et al., 2015; Hänsch and Mendel, 2009; Palmer and Guerinot, 2009).

Improving plant iron content is of primary importance because iron deficiency also affects billions of people worldwide (World Health Organization, https://www.who.int) and because plants serve as the main source of dietary iron for most of the world's population. The primary control point for plant iron homeostasis is iron uptake in root epidermal cells. In the model plant *Arabidopsis thaliana*, and more generally in non-grasses plants, this crucial step involves the high affinity iron uptake complex sitting at the plasma membrane and composed of the AHA2 H^+^-ATPase, the FRO2 ferric chelate reductase and the iron transporter IRT1. The proton excretion mediated by AHA2 increases ferric iron solubility that will be then reduced into ferrous iron by the FRO2 reductase before being taken up in root epidermal cells by IRT1 (Jeong et al., 2017; Martin-Barranco et al., 2020; Palmer and Guerinot, 2009; Thomine and Vert, 2013). Defects in acidification, iron reduction or transport, as observed in *aha2*, the *FRO2* gene mutant *frd1*, or *irt1* or mutants in upstream regulators are all associated with reduced plant growth and chlorosis pointing to their crucial role in iron uptake soil (Santi and Schmidt, 2009; Vert et al., 2002; Yi and Guerinot, 1996). Consequently, the expression of the corresponding iron uptake machinery genes are rapidly and strongly upregulated upon low iron conditions to increase iron absorption from the soil (Robinson et al., 1999; Santi and Schmidt, 2009; Vert et al., 2002). Besides, plants use an intricate signaling network where various hormonal or stress pathways impinge on iron deficiency signaling to adjust iron uptake to plant growth (Gao and Dubos, 2021; Hanikenne et al., 2021; Spielmann and Vert, 2021).

IRT1 belongs to the ZIP gene family, whose members have been implicated in the transport of a variety of divalent metals. IRT1 is now seen as a broad spectrum metal transporter, capable of transporting Fe, zinc (Zn) and manganese (Mn) with high affinity kinetics. Plant grown in iron-deficient conditions and strongly expressing *IRT1* tend to accumulate Zn or Mn, which can have detrimental effects if present at too high levels in plant cells (Vert et al., 2002). Plants have evolved a very sophisticated strategy to limit the overaccumulation of Zn and Mn driven by IRT1. IRT1 was shown to act as a bifunctional transporter-receptor, capable of sensing Zn or Mn ion fluxes using a histidine-rich motif in a cytosolic regulatory loop (Cointry and Vert, 2019; Dubeaux et al., 2018). Zn and Mn bind directly to such histidine residues when plants face excess of these metals, triggering the recruitment of the CIPK23 kinase. CIPK23 phosphorylates serine and threonine residues in IRT1 regulatory loop, thus creating a docking site for the IDF1 RING E3 ubiquitin ligase that in turn drives the lysine(K) 63 polyubiquitination of IRT1 (Dubeaux et al., 2018). Ubiquitinated IRT1 is then sorted to the vacuole and degraded to limit the accumulation of highly reactive and potentially toxic metals when found in excess (Barberon et al., 2011; Dubeaux et al., 2018).

The fact that an excess of non-iron metal substrates of IRT1 such as Zn or Mn leads to the removal of IRT1 from the cells surface and its degradation decreases the uptake of corresponding metals. In particular, this may limit the efficiency of IRT1-based strategies to increase the uptake of such metals in biofortification or phytoremediation approaches. Besides Zn or Mn, IRT1 likely transports other metals including cadmium (Cd) (Barberon et al., 2011; Connolly et al., 2002; Eide et al., 1996; Vert et al., 2002). IRT1 expression in wild-type yeast indeed confers hypersensitivity to external cadmium (Cd), and that Cd competes with Fe uptake by IRT1 when expressed in the iron transport-defective *fet3fet4* yeast mutant strongly suggests that IRT1 has the ability to transport Cd (Barberon et al., 2011; Eide et al., 1996). IRT1 also appears to transport Cd in plants as i) the *irt1* mutant fails to accumulate Cd in response to low Fe conditions, and ii) plants overexpressing IRT1 are hypersensitive to Cd and accumulate Cd in their roots (Connolly et al., 2002; Vert et al., 2002). Hence, IRT1 is considered as the major entry route for Cd in Fe-deficient plants and in the food chain. As a consequence, IRT1 represent a promising target to engineer plants with increased Cd uptake and storage capacities to depollute or rehabilitate polluted sites such as industrial plants or mines with high Cd levels. Here we addressed whether IRT1 is post-translationally regulated by Cd and the underlying mechanisms. We uncovered that Cd excess triggers IRT1 internalization and degradation, albeit with a lower efficiency than for Zn. The Cd-mediated endocytosis of IRT1 involves direct binding of Cd to histidine residues within IRT1 regulatory loop and IDF1-dependent ubiquitination of two lysines. Altogether our work shed light on the post-translational control of IRT1 by Cd that may limit the use of IRT1 in phytoremediation strategies.

## Results

### IRT1 is endocytosed in response to cadmium

The broad spectrum root metal transporter IRT1 has previously been shown to undergo internalization and targeting to the lytic vacuole for degradation when plants are exposed to an excess of its secondary metal substrates zinc, manganese or cobalt (Dubeaux et al., 2018). Considering that Cd is also a substrate of IRT1 when expressed in yeast or in plants (Barberon et al., 2011; Eide et al., 1996; Rogers et al., 2000; Vert et al., 2001; Vert et al., 2002), we wondered whether Cd excess also triggers the degradation of IRT1 by ubiquitin-mediated endocytosis. To address this, we took advantage of a reporter line expressing the functional IRT1-mCitrine fusion (IRT1-mCit) (Dubeaux et al., 2018). Similar to other non-iron metal substrates of IRT1 (Dubeaux et al., 2018), Cd likely affects *IRT1* transcription by competing with Fe for uptake (Fig. S1A). We therefore generated a PIN2::IRT1-mCit to constitutively express the fluorescent IRT1 reporter line at the root tip, where imaging of endocytosis is most appropriate due to the reduced cell size and best characterized. PIN2::IRT1-mCit plants were grown in the absence of iron and non-iron metals (Zn, Mn and Co), condition when *IRT1* is normally expressed, and exposed or not to short term metal excess. The PIN2 promoter yielded constitutive IRT1-mCit mRNA accumulation, irrespective of Cd or other non-iron metal substrates of IRT1 (Fig. S1B). Under standard growth conditions, IRT1-mCit was observed at the plasma membrane and in endosomal compartments, consistent with published observations (Barberon et al., 2014; Barberon et al., 2011; Dubeaux et al., 2018) (Fig. 1A). An excess of Zn, Mn and Co strongly decreased IRT1-Cit levels at the plasma membrane and concomitantly increased its endosomal accumulation, as previously reported (Dubeaux et al., 2018) (Fig. 1A). Cd also triggered IRT1-mCit endocytosis highlighted by a reduced plasma membrane localization of IRT1 and an increased number of endosomes, albeit to a lesser extent than what is observed for the other non-iron metal substrates of IRT1 (Fig. 1A). Quantification of the IRT1-mCit levels at the cell surface clearly showed that Cd treatment impacts on the localization of IRT1 at the plasma membrane (Fig. 1B). To better assess the influence of metals on IRT1 distribution and limit the impact of variation in fluorescence between images or after metal treatment, we also quantified the ratio between plasma membrane and intracellular pools of IRT1-mCit. As expected, a severe drop in such ratio was observed upon Zn, Mn and Co excess (Fig. 1C). A short term excess of Cd (50μM) also translated into a decreased ratio between plasma membrane and intracellular IRT1-mCit (Fig. 1C). However, Cd appeared again less potent at the concentration used than the combined excess of Zn, Mn and Co to provoke IRT1 endocytosis. We therefore increased Cd concentrations to address whether IRT1-mCit endocytosis can reach the same extent as with other non-iron metal substrates of IRT1. However, the extreme toxicity of Cd prevented us from reaching a clear conclusion at higher cadmium concentration (75μM Cd) due to presence of dead cells highlighted by intracellular propidium iodide accumulation (Fig S2A). Moreover, high cadmium concentration also perturbed plasma membrane level of the brassinosteroid receptor BRI1 that is not related to metal homeostasis (Fig. S2B and S2C). We therefore used a concentration of Cd of 50 μM to monitor IRT1-mCit protein dynamics in the rest of the manuscript.

**Figure 1:**
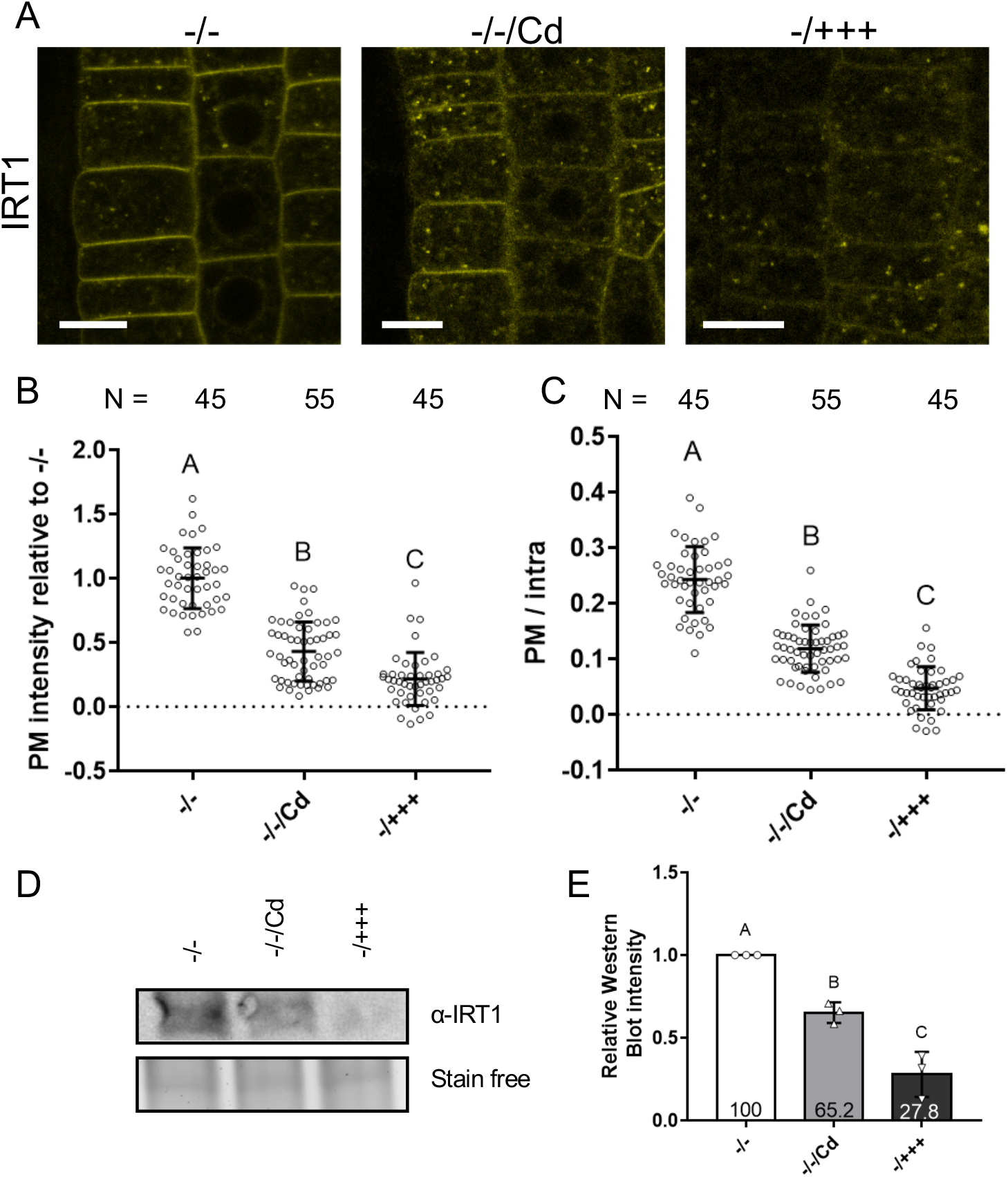
Cadmium regulates IRT1 endocytosis and degradation. (A) Confocal microscopy analyses of root epidermal cells from plants expressing PIN2::IRT1-mCit (IRT1). Plants were grown in medium without iron and without non-iron metals (Zn, Mn and Co), then liquid-treated for 24h with the same medium (−/−), with 50μM cadmium (−/−/Cd) or with an excess of non-iron metals (−/+++). Scale bars, 10μm. Quantification of plasma membrane (PM) signal intensity (B) or ratio of plasma membrane to intracellular signal (C) of IRT1-mCit from experiments performed in (A). Error bars represent SD and number of values are indicated in the graph. Different letters indicate significant differences between condition (one-way ANOVA, Tukey post-test, *** p < 0.001). Western blot analyses (D) and protein quantification (E) of plants expressing 35S::IRT1 and grown as in (A). Protein levels were detected using anti-IRT1 antibodies on root protein extracts. Detection of total proteins using stain free technology is used as loading control. (E) Value are means of 3 biological replicate and error bars represent SD. Different letters indicate significant differences between condition (one-way ANOVA, Tukey post-test, *** p < 0.001).

To better examine the influence of Cd on IRT1 protein accumulation, we performed western blot analyses using the 35S::IRT1 transgenic line constitutively expressing *IRT1* and anti-IRT1 antibodies (Barberon et al., 2011). Using root extracts from plants grown under standard conditions and then treated or not with metals excess, or 50μM Cd, we reproducibly observed reduced IRT1 protein levels for plants exposed to Cd or combined Zn, Mn and Co excess (Fig. 1D). Quantification of signal intensities confirmed the decrease of IRT1-mCit protein accumulation of around 35% and 72%, respectively, after cadmium exposure or combined Zn, Mn and Co excess (Fig. 1E), mediated at least in part by a Cd-dependent post-translational regulation triggering IRT1 endocytosis.

Altogether, these observations indicate that Cd excess results in IRT1 endocytosis and degradation, likely to limit Cd overaccumulation in plant tissues.

### Cadmium is directly sensed by the histidine-rich motif in IRT1

IRT1 was shown to be a bifunctional transporter-receptor (Cointry and Vert, 2019; Dubeaux et al., 2018), capable of sensing non-iron metals using a histidine-rich stretch sitting in the intracellular regulatory loop to control its own levels by ubiquitin-mediated endocytosis and degradation. The expression of an IRT1_4HA_ mutant version substituted for the four histidine residues involved in non-iron metal binding indeed prevents IRT1 degradation upon non-iron metal excess (Dubeaux et al., 2018). Since Cd is also promoting IRT1 endocytosis, we evaluated if such post-translational regulation also uses direct Cd binding to histidine residues in IRT1 regulatory loop by studying the response to Cd of the IRT1_4HA_-mCit reporter line. In contrast to the PIN2::IRT1-mCit line that showed decreased plasma membrane levels and increased endosomal levels of IRT1-mCit upon Cd exposure (Fig. 2A), the PIN2::IRT1_4HA_-mCit transgenic line harbored slightly higher levels at the plasma membrane and no visible change in intracellular IRT1-mCit levels after Cd treatment (Fig. 2A). Quantification of the plasma membrane fluorescence intensity confirmed the limited impact of Cd on IRT1_4HA_ endocytosis compered to wild-type IRT1-mCit (Fig. 2B). Surprisingly, the effect of Cd on plasma membrane to intracellular ratio of fluorescence for IRT1_4HA_ was comparable to that of wild-type IRT1mCit-expressing plants (Fig. 2C). Since IRT1_4HA_-mCit shows a lot more endosomes than IRT1-mCit even in standard conditions (Dubeaux et al., 2018), this likely masks the influence of Cd on IRT1-mCit endocytosis.

**Figure 2:**
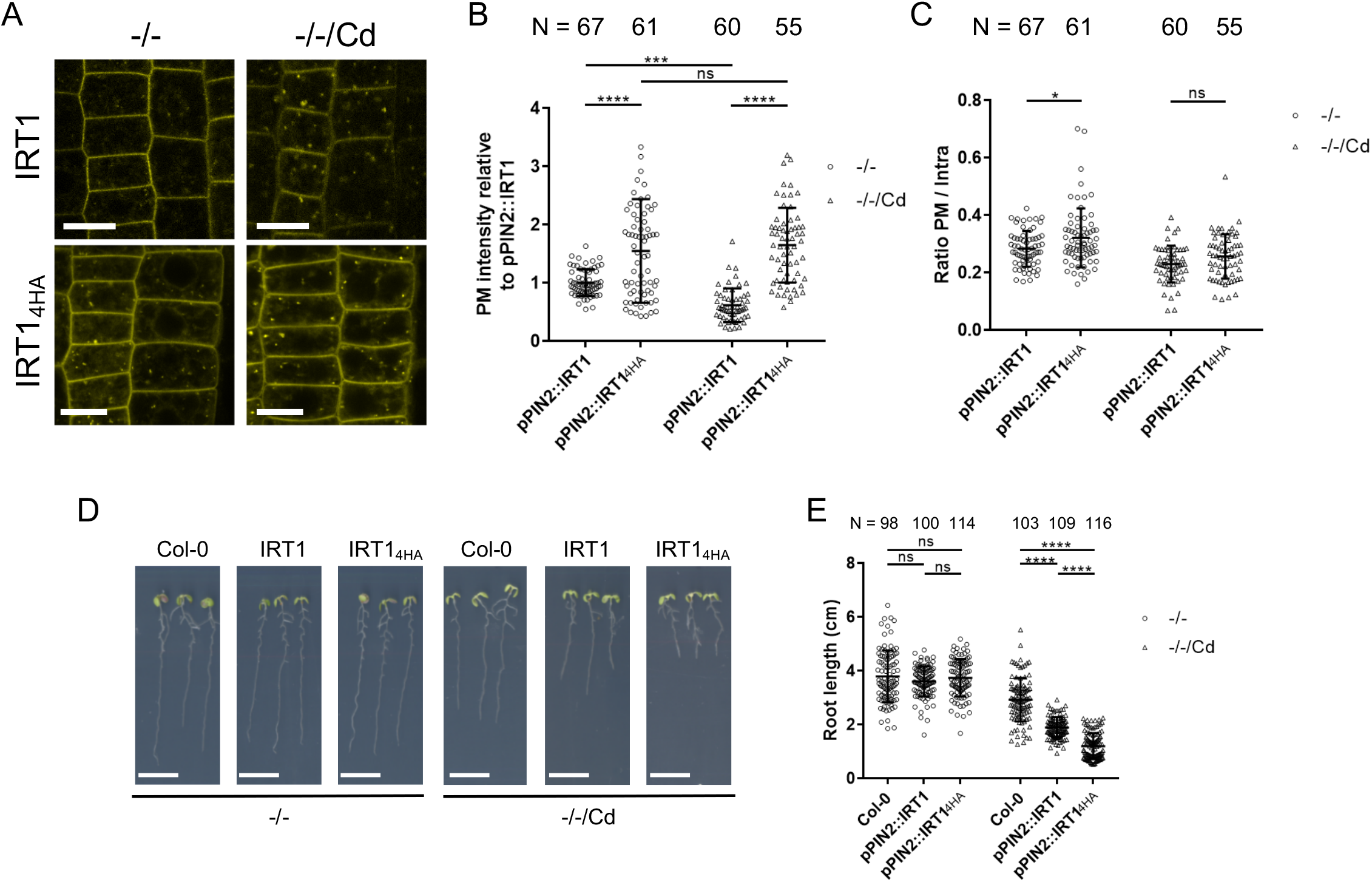
The histidine-rich motif of IRT1 is essential for cadmium-mediated endocytosis. (A) Confocal microscopy analyses of root epidermal cells from plants expressing PIN2::IRT1-mCit (IRT1) or PIN2::IRT1_4HA_-mCit (IRT1_4HA_). Plant were expose to cadmium (−/−/Cd) or not (−/−) for 24h. Scale bars, 10μm. Quantification of plasma membrane (PM) signal intensity (B) or ratio of plasma membrane to intracellular signal (C) of IRT1-mCit or IRT1_4HA_-mCit from experiments performed in (A). Error bars represent SD and number of values are indicated in the graph. Asterisk indicate significant (Two-way ANOVA, Sidak post-test, **** p < 0.0001; ** p < 0.01; * p < 0.05; ns, Not significant). Phenotype (D) and root length measurement (E) of 14-day-old wild-type (Col-0), plant expressing PIN2::IRT1-mCit (IRT1) or PIN2::IRT1_4HA_-mCit (IRT1_4HA_) germinated and grown in medium without iron and without non-iron metals (Zn, Mn and Co) (−/−) or the same medium supplemented with cadmium (−/−/Cd). Error bars represent SD and number of values are indicated in the graph. Asterisk indicate significant differences between condition (Two-way ANOVA, Sidak post-test, **** p < 0.0001; ns, not significant). Scale bars, 1 cm.

To better examine the role of the IRT1 histidine-rich stretch in plant expose to cadmium, we grew wild-type, PIN2::IRT1-mCit and PIN2:: IRT1_4HA_-mCit plants on control or Cd-containing media and measured root length. No difference were observed in control condition (Fig. 2D, E). When exposed to Cd, PIN2::IRT1-mCit was slightly more sensitive to Cd than wild-type, indicating that the expression of IRT1-mCitrine in the sole meristem is sufficient to generate hypersensivity to external Cd. Expression of IRT1_4HA_-mCit however yielded a strong sensitivity to Cd as visualized by the reduced root length upon Cd exposure (Fig. 2D, E), and by the swelling of the root meristem (Fig. S3). This is consistent with an increased Cd uptake due to the constitutive expression of IRT1 and due to the higher amount of IRT1_4HA_ at the plasma membrane. Overall, this indicates that the histidine-rich stretch is required for IRT1 to properly respond to Cd.

At subtoxic concentrations, Cd appeared less effective than Zn, Mn and Co in triggering IRT1 endocytosis. This may be explained by a reduced affinity of the histidine-rich motif in IRT1 regulatory loop for Cd. To confirm this hypothesis, we implemented microscale thermophoresis to directly monitor the binding and dissociation constant (Kd) of metals to the regulatory histidine residues in IRT1. Purified peptides encompassing the regulatory loops of wild-type IRT1 and its IRT1_4HA_ counterpart were expressed and purified from *Escherichia coli*. Peptides were labelled and incubated with decreasing concentrations of Zn or Cd before temperature-induced variations in IRT1 peptide fluorescence were recorded. Consistent with previously reported using other approaches (Dubeaux et al., 2018), Zn bound to the regulatory loop of IRT1 (Fig. 3A). Such binding was abolished when the four histidine residues found in the histidine-rich stretch were substituted, confirming that Zn directly binds to histidines in the regulatory loop of IRT1. Analyses of the microscale thermophoresis spectra yielded a Kd in the micromolar range (19.5 ± 6.3μM) for Zn for the IRT1 wild-type peptide (Fig. 3B). Cd also bound to the wild-type IRT1 peptide in microscale thermophoresis experiments, but with a Kd in the millimolar range (2.5 ± 0.6mM) (i.e. 2 orders of magnitude higher) (Fig. 3A, B). Similar to what was observed for Zn, substitution of the four histidine residues in IRT1 regulatory loop completely abolished Cd binding.

**Figure 3:**
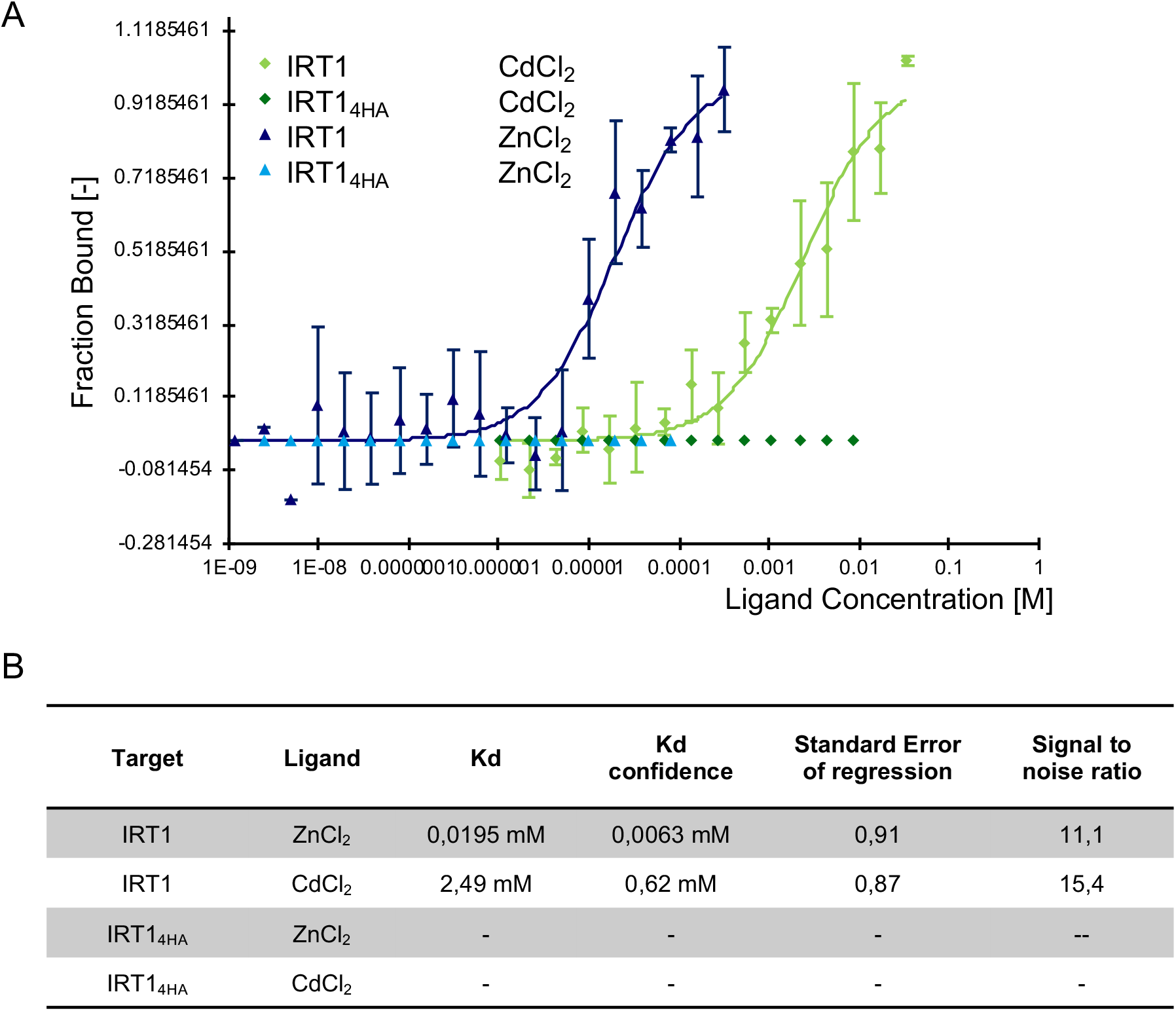
IRT1 loop directly binds zinc and cadmium using its histidine-rich stretch. (A) Microscale thermophoresis analyses of wild-type IRT1 regulatory loop (IRT1) or variant mutated for the histidine-rich motif (IRT1_4HA_) with zinc or cadmium. Dark blue triangles, IRT1 with zinc; light green diamonds, IRT1 with cadmium; light blue triangles, IRT1_4HA_ with zinc; dark green diamonds, IRT1_4HA_ with cadmium). The mutated peptide for the histidine-rich motif (IRT1_4HA_) failed to fit to a binding curve with both zinc and cadmium. Value are means of biological duplicates with at least two technical replicates for each and error bars represent SD. (B) Kd determination using curve presented in (A).

Taken together, these results confirm the ability of IRT1 to sense transported Cd levels using its histidine-rich stretch to regulate its own stability. The fact that IRT1 efficiently transports Cd, but shows lower affinity to sense Cd compared to other non-iron metal substrates, contributes to the extreme lethality observed for Cd.

### Cadmium-mediated endocytosis of IRT1 is mediated by ubiquitination

We next investigated the molecular mechanisms driving the Cd-mediated endocytosis of IRT1. Considering that IRT1 undergoes ubiquitin-mediated endocytosis and vacuolar degradation upon Zn, Mn and Co excess (Dubeaux et al., 2018), we speculated that elevated Cd concentrations may use the same mechanism and machinery. Two lysine residues in the regulatory loop of IRT1, K154 and K179, are required for IRT1 endocytosis in standard and Zn, Mn and Co excess conditions (Barberon et al., 2014; Barberon et al., 2011; Dubeaux et al., 2018). We therefore generated a PIN2::IRT1_2KR_-mCit transgenic line where both lysine residues are substituted to arginines, to maintain the positive charge but render IRT1 non-ubiquitinatable. IRT1_2KR_-mCit-expressing plants showed a strong plasma membrane localization of IRT1 with little or no endosomes (Fig. 4A), as expected. In contrast to what is observed for the wild-type IRT1-mCit where endosomes positive for IRT1-mCit were observed after Cd exposure, IRT1_2KR_-mCit-expressing plants showed no sensitivity to Cd regarding IRT1 endocytosis (Fig. 4A). Indeed, IRT1_2KR_-mCit was almost never observed in endosomes even upon Cd exposure, clearly pointing to the role of lysine residue K159 and K174 for IRT1 endocytosis in the response to Cd (Fig. 4A). Quantification of the plasma membrane to intracellular fluorescence ratio for IRT1_2KR_-mCit however revealed no difference of sensitivity upon Cd treatment with IRT1-mCit (Fig. 4C). This is likely a consequence of enhanced cadmium toxicity observed in IRT1_2KR_-mCit-expressing plants. We indeed noticed that Cd led to a decrease of plasma membrane levels of IRT1_2KR_-mCit in the absence of a concomitant increase in endosomes, reminiscent of what we observed with BRI1 on higher cadmium concentration (Fig. 4A, 4B and Fig. S1). Also, dying cells were observed for IRT1_2KR_-mCit-expressing plants at 50μM Cd (Fig. S4A). Besides, to ascertain that the lack of IRT1_2KR_-mCit-positive endosomes observed when plants are challenged with Cd do not result from a general defect in endocytosis, we analyzed FM4-64 internalization after cadmium exposure. No difference in FM4-64 positive endosomes could be observed between IRT1-mCit and IRT1_2KR_-mCit at 50μM Cd (Fig. S4B). Altogether, this indicates Cd specifically triggers IRT1 endocytosis and that residues K154 and K179 are required for the Cd-mediated internalization of IRT1.

**Figure 4:**
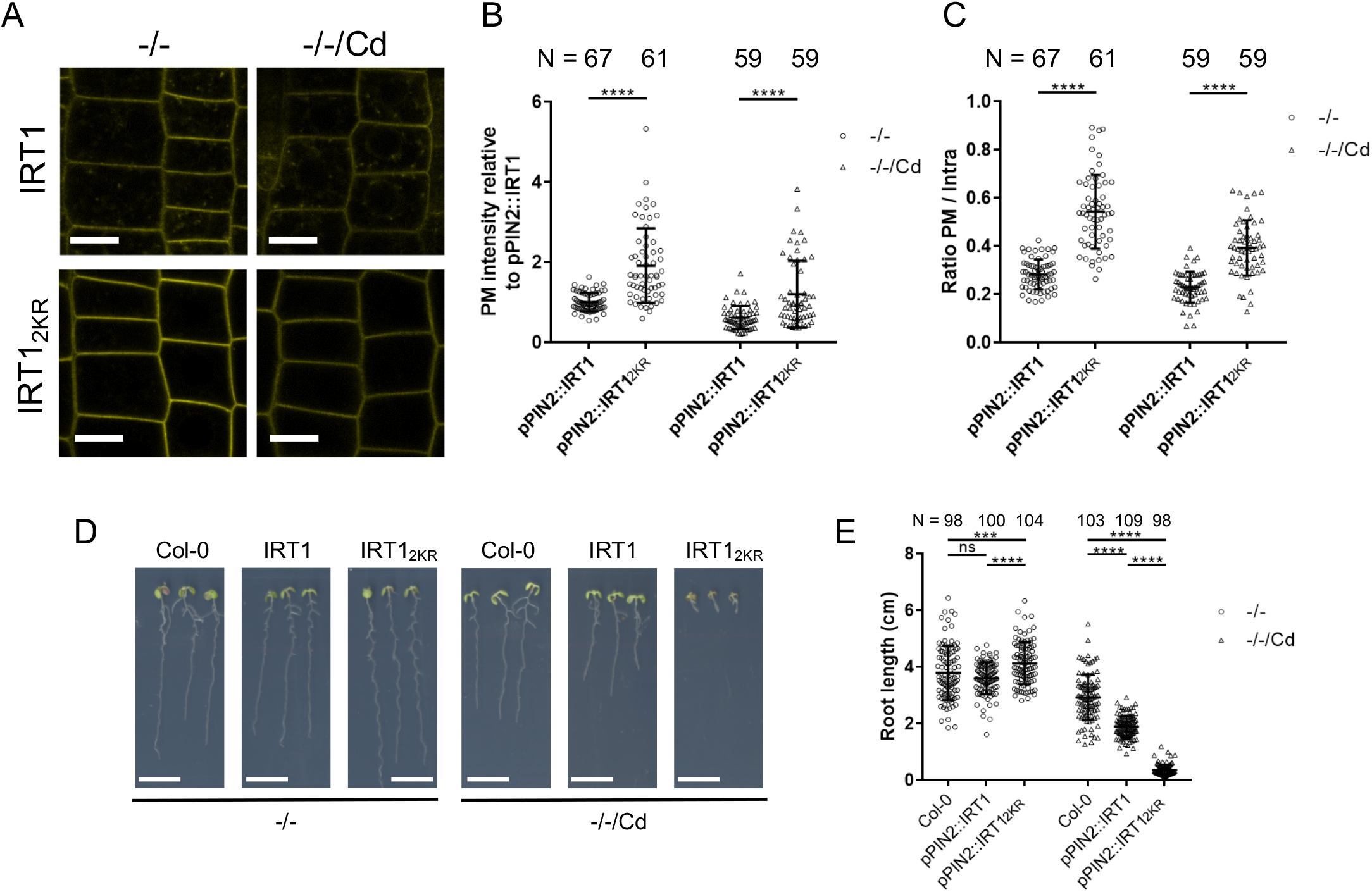
Lysine residues K154 and K179 are crucial for IRT1 cadmium-induced endocytosis. (A) Confocal microscopy analyses of root epidermal cells from plants expressing PIN2::IRT1-mCit (IRT1) or PIN2::IRT1_2KR_-mCit (IRT1_2KR_). Plant were exposed to cadmium (−/−/Cd) or not (−/−) for 24h. Scale bars, 10μm. Quantification of plasma membrane (PM) signal intensity (B) or ratio of plasma membrane to intracellular signal (C) of IRT1-mCit or IRT1_2KR_-mCit from experiments performed in (A). Error bars represent SD and number of values are indicated in the graph. Asterisk indicate significant differences between genotype (Two-way ANOVA, Sidak post-test, **** p < 0.0001). PIN2::IRT1-mCit values used in panel (B) and (C) are identical for those presented in Figure 2A and 2B. Phenotype (D) and root length measurement (E) of 14-day-old wild-type (Col-0), plant expressing PIN2::IRT1-mCit (IRT1) or PIN2::IRT1_2KR_-mCit (IRT1_2KR_) germinated and grown in medium without iron and without non-iron metals (Zn, Mn and Co) (−/−) or in the same medium with cadmium (−/−/Cd).. Error bars represent SD and number of values are indicated in the graph. Asterisk indicate significant differences between condition (Two-way ANOVA, Sidak post-test, **** p < 0.0001; *** p < 0.001; ns, not significant). Scale bars, 1cm.

We next examined the impact of the loss of IRT1 ubiquitination at residues K154 and K179 would impact plant growth responses to Cd exposure. We therefore grew wild-type, PIN2::IRT1-mCit and PIN2:: IRT1_2KR_-mCit plants on control or Cd-containing media and measured root length. PIN2::IRT1_2KR_-mCit plants displayed slightly longer roots in control conditions, likely due to their increased ability to take up traces of iron (Fig. 4D, 4E). However, when grown in the presence of Cd, IRT1_2KR_-mCit-expressing plants became dramatically short, consistent with the absence of Cd-mediated endocytosis in this background. It is noteworthy that the expression of IRT1_2KR_-mCit in the epidermis and cortical cells of the meristematic zone is sufficient to generate detrimental growth defects by Cd. This is explained by the strong alteration of the meristem in IRT1_2KR_-mCit plants observed in the presence of Cd (Fig. S3), as visualized by propidium iodide staining, further attesting of the uncontrolled Cd uptake when IRT1 ubiquitination is prevented.

### IDF1 controls the cadmium-induced endocytosis of IRT1

The ubiquitination of IRT1 involves the IDF1 RING E3 ubiquitin ligase that covalently attaches ubiquitin moieties to lysine residues K159 and K179 (Dubeaux et al., 2018). Since the endocytosis of IRT1 observed upon Cd excess requires both lysine residues, we tested the possible involvement of IDF1 in this process. To do so, we crossed the PIN2::IRT1-mCit line to *idf1* mutants and obtained the double homozygous progeny. We next monitored the influence of Cd treatment on the distribution of IRT1-mCit in the wild-type or *idf1* mutant background. A short term Cd exposure yielded a low ratio between the plasma membrane and intracellular pools of IRT1-mCit when expressed in the wild-type background (Fig. 5A, 5C). In the *idf1* mutant however, fewer endosomes were observed and plasma membrane level do not decreased after Cd treatment (Fig. 5A, 5B), leading to an increase in the abovementioned ratio (Fig. 5C), pointing to the requirement for IDF1 to destabilize IRT1 when plants face Cd. Consistent with the observation that IDF1 is required for the Cd-mediated endocytosis of IRT1, *idf1* mutant is hypersensitivity to Cd compared to wild-type (Fig. 5D, 5E).

**Figure 5:**
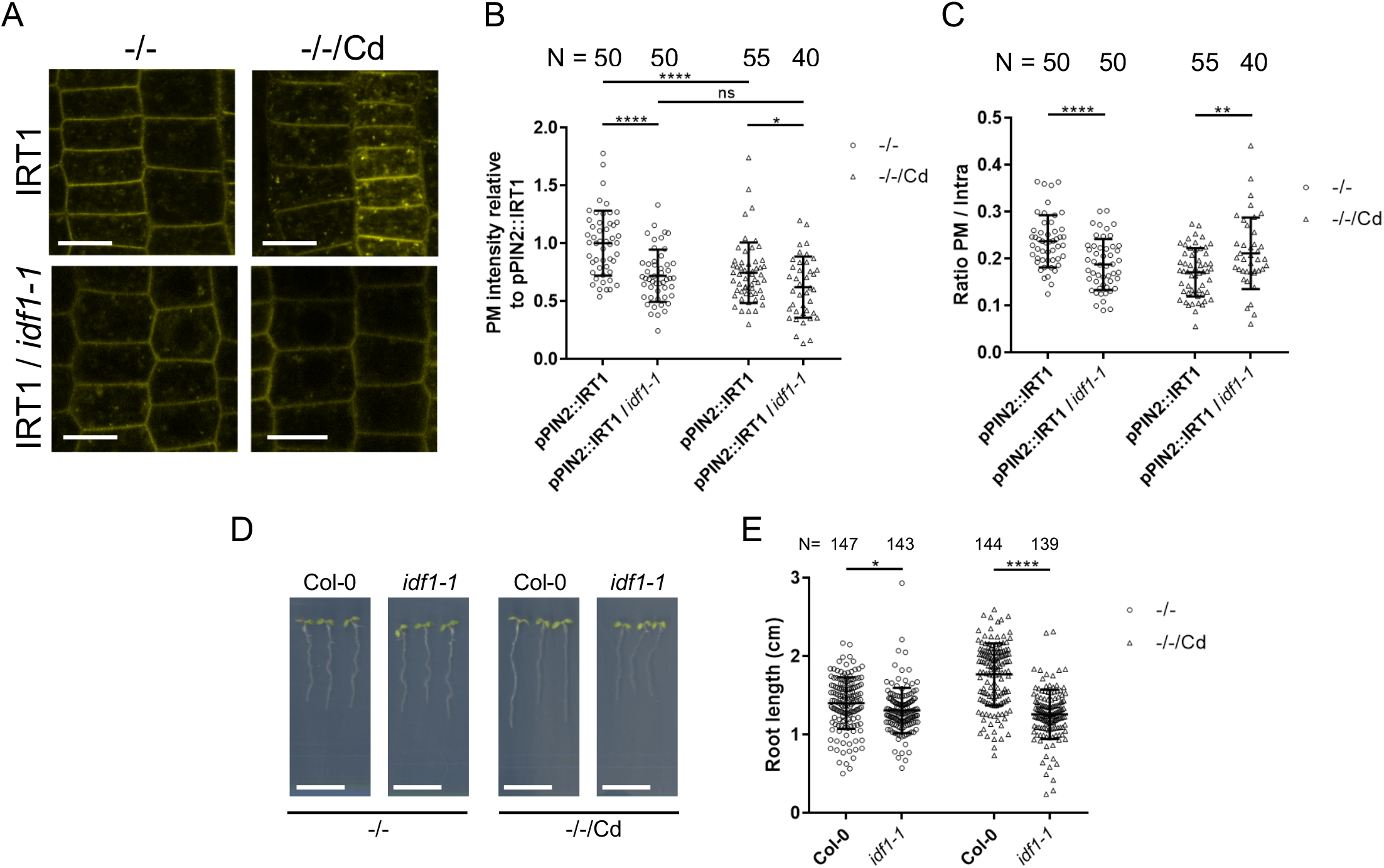
IDF1 is involved in IRT1 endocytosis in response to cadmium. (A) Confocal microscopy analyses of root epidermal cells from plants expressing PIN2::IRT1-mCit (IRT1) in wild-type or *idf1-1* background (*idf1-1*/IRT1). Plant were exposed to cadmium (−/−/Cd) or not (−/−) for 24h. Scale bars, 10μm. Quantification of plasma membrane (PM) signal intensity (B) or ratio of plasma membrane to intracellular signal (C) of IRT1-mCit in wild-type or *idf1-1* background. Error bars represent SD and number of values are indicated in the graph. Asterisk indicate significant differences between genotype (Two-way ANOVA, Sidak post-test, **** p < 0.0001; ** p < 0.01; ns, not significant). Phenotype (D) and root length measurement (E) of 7-day-old wild-type or *idf1-1* plants germinated and grown in medium without iron and without non-iron metals (Zn, Mn and Co) (−/−) or the same medium with cadmium (−/−/Cd). Error bars represent SD and number of values are indicated in the graph. Asterisk indicate significant differences between condition (Two-way ANOVA, Sidak post-test, **** p < 0.0001; *** p < 0.001; ns, not significant). Scale bars, 1cm.

Taken together, these results clearly argue for a role of IDF1-mediated ubiquitination of lysine residues K159 and K174 when plants are challenged with Cd, and provide a deeper understanding about the molecular mechanisms driving IRT1 endocytosis when plants face Cd.

## Discussion

The root iron transporter IRT1 shows a broad metal transport spectrum by transporting not only ferrous iron, but also zinc, manganese, cobalt and cadmium (Barberon et al., 2011; Eide et al., 1996; Vert et al., 2002). Iron is the primary substrate of IRT1, since *IRT1* transcription is rapidly and strongly induced upon low iron conditions and the severe phenotype displayed by the *irt1* loss-of-function mutant is reversed by exogenous application of iron only (Vert et al., 2002). Zinc, manganese and cobalt, which accumulate in plant tissues upon low iron conditions, rather control IRT1 post-translationally leading to its ubiquitin-mediated endocytosis and degradation (Barberon et al., 2011; Dubeaux et al., 2018). Such removal of IRT1 from the cell surface allows to limit the accumulation of highly reactive non-iron metals transported by IRT1. Inability of IRT1 to respond to zinc, manganese or cobalt excess leads to a dramatic accumulation of the corresponding metals in plant tissues and plant death (Barberon et al., 2011). IRT1 also represents the major entry route for cadmium in plants under iron-limited conditions (Connolly et al., 2002; Vert et al., 2002). Here, we have demonstrated that, similarly to the other non-iron metal substrates of IRT1, cadmium excess is directly sensed by the histidine residues in the IRT1 regulatory loop to trigger its endocytosis and degradation.

Cadmium is an important environmental pollutant emanating from anthropogenic activities such power plants or waste incinerators (Sanità di Toppi and Gabbrielli, 1999). Plants take up cadmium and are the entry route cadmium into the food chain, where it gradually accumulates and raises a serious threat to human health. Understanding the precise molecular mechanisms underlying cadmium transport and responses to cadmium excess in plants in therefore crucial. In our conditions, sub-lethal concentrations of cadmium yield regulate *IRT1* transcriptionally and post-translationally. The accumulation of *IRT1* mRNA observed upon cadmium excess is consistent with cadmium competing with iron for transport, thus boosting *IRT1* transcription (Fig. S1A). This is also in accordance with what has been observed for a combined excess of the other non-iron metal substrates of IRT1 (Dubeaux et al., 2018). *IRT1* mRNA accumulation was previously reported to be negatively impacted by cadmium, leading to a decrease in IRT1 protein (Connolly et al., 2002). Such discrepancy may be explained by the much higher concentration used in the latter study that may cause cell death and global downregulation of gene expression compared to our conditions.

The cadmium-induced endocytosis of IRT1 appears to use the same mechanisms reported for other non-iron metal substrates of IRT1. Cadmium is directly sensed by the histidine-rich motif found in the regulatory loop of IRT1 (Fig. 3). Mutation of such histidine residues impairs IRT1 endocytosis upon cadmium excess and consequently yields strong hypersensitivity to cadmium (Fig. 2). The removal of IRT1 from the cell surface and its degradation likely uses ubiquitin-mediated endocytosis since it is abolished for the ubiquitination-defective IRT1_2KR_ mutant or in the *idf1* mutant background (Fig. 4 and 5). Both show hypersensitivity to cadmium excess, further supporting their functional importance for IRT1 to properly respond to cadmium (Fig. 4 and 5). However, at the concentrations of 50μM used in this study, cadmium appears to be less potent than the combined excess of zinc, manganese and cadmium since yielding lower endocytosis and degradation (Fig. 1). Since slightly higher concentrations of cadmium are associated with cell death (Fig. S2), this suggests that cadmium sensing by IRT1 is not as efficient as for its other non-iron metal substrates. We confirmed this hypothesis by measuring a dissociation constant of cadmium for the metal sensing histidine-rich domain of IRT1 two orders of magnitude lower than the one for zinc (Fig. 3). Such reduced ability to degrade IRT1, combined to cadmium being efficiently transported by IRT1 across the plasma membrane, contributes to the extreme toxicity and lethality of cadmium in plants.

The phytoremediation- or phytoextraction-based processes of using plants to rehabilitate cadmium-contaminated soils or mine for cadmium emerge as an eco-friendly and inexpensive strategies. Only a few cadmium hyperaccumulating plants have been identified to date (Krämer, 2010; Meyer and Verbruggen, 2012; Reeves et al., 2018), due to the non-essential nature of this metal. Despite the ability of cadmium hyperaccumulators to accumulate more than 0.01% of cadmium in their shoot dry biomass, they often display reduce stature that greatly limits their use in phytoremediation or phytoextraction. As the primary actor responsible for cadmium uptake in plants, IRT1 stands out as a major target to engineer plants with enhanced cadmium uptake capacities in and a robust root system to explore the soil. This can in theory be achieved by increasing the cadmium uptake specific activity of IRT1 and/or by altering its cadmium-induced endocytosis. Substitution of residues in IRT1 allowed the creation of variants with altered metal selectivity when expressed in yeast (Rogers et al., 2000), but the kinetics of metal transport were not addressed. Recently, the first tridimensional structure of a bacterial ZIP transporter was released (Zhang et al., 2017), which together with the Alphafold-based model prediction for IRT1 (Jumper et al., 2021), may help scientists to generate an IRT1 variant with increased cadmium transport properties. The second approach is to decrease further the cadmium-mediated endocytosis of IRT1. Although IRT1 is much less sensitive to cadmium than to its other non-iron metal substrates for metal-regulated endocytosis (Fig. 1), we have clearly shown that mutating the histidine residues in the regulatory loop of IRT1 abolishes IRT1 degradation and confer strong hypersensitivity to cadmium (Fig. 2). The sole approach of limiting IRT1 endocytosis would result in a non-specific increase in metal transport, pointing to the need of a mix the two strategies. To this purpose, detailed structural analyses of IRT1 in the presence/absence of its various metal substrates is still awaited.

Boosting metal transport by expression of an engineered IRT1 must be accompanied with increased cadmium root-to-shoot translocation and storage/detoxication capacities to ensure maximum tolerance to high metal content. Extensive efforts over the past two decades to characterize the molecular determinants of cadmium homeostasis now provide us with some possible strategies to be combined to highly stable and highly selective boosted versions of IRT1 to engineer artificial cadmium hyperaccumulating plants. *HMA2* and *HMA4* genes encode plasma membrane P-type ATPase responsible for zinc and cadmium loading into the xylem and long distance transport to shoots (Hussain et al., 2004; Sinclair et al., 2018; Wong and Cobbett, 2009). Enhanced *HMA4* expression in *A. thaliana* increased root cadmium tolerance and shoot sensitivity (Hanikenne et al., 2008; Nouet et al., 2015). Consistently, the downregulation of *A. halleri HMA4* yields a strong reduction in cadmium accumulation in shoots (Hanikenne et al., 2008). Similarly, interfering the expression of *A. halleri ZIP6*, a zinc and cadmium cytoplasmic influx pump, differentially impact cadmium tolerance in *A. thaliana* and *A. halleri* (Spielmann et al., 2020). The nicotianamine metal chelator encoding gene *NAS2* also appears as an interesting target to alter cadmium translocation since downregulation of *A. halleri NAS2* leads to decreased shoot cadmium levels (Deinlein et al., 2012). The sequestration of cadmium into vacuoles uses the tonoplast P-type ATPase HMA3 in non-hyperaccumulating plants such as rice and hyperaccumulators (Fischer et al., 2017; Liu et al., 2017; Miyadate et al., 2011; Ueno et al., 2010). The combination of some of these actors will be necessary to generate an artificial hyperaccumulator better suited to phytoremediate cadmium polluted soils.

## Methods

### Constructs and generation of transgenic plants

The pDONR.P4P1R-PIN2, pDONR221-IRT1-mCit, IRT1-mCit variants and pDONR.P2RP3-Mock Gateway entry clones were previously described (Dubeaux et al., 2018; Marquès-Bueno et al., 2016). Final destination PIN2::IRT1-mCit construct and variants (PIN2::IRT1_2KR_-mCit, PIN2::IRT1_4HA_-mCit) were obtained using multisite Gateway recombination system (Life Technologies) using the pG7m34GW destination vector, pDONR.P4P1R-pPIN2, pDONR221-IRT1-mCit (or variant) and pDONR.P2RP3-Mock entry clones. Col-0 plant were transformed by the floral dipping method using *Agrobacterium tumefaciens*. Around 20 independent T1 lines were isolated and three representative mono-insertion lines were selected in T2. Independent lines homozygous for the transgene were selected in T3.

### Plant material and growth conditions

*Arabidopsis thaliana* (accession Columbia-0 (Col-0)), PIN2::IRT1-mCitrine and variant (generated in this study), p35S::IRT1 (Barberon et al., 2011) and *idf1-1* mutant (Dubeaux et al., 2018) were used. *idf1-1* mutant line was crossed with pPIN2::IRT1-mCitrine to obtain *idf1-1*/PIN2::IRT1-mCitrine. After an ethanol sterilization, seeds were grown on solid agar (1% w/v; New Kalys agar; Kalys) half-strength Murashige and Skoog medium lacking iron and non-iron metals (-Fe/-Metals; −/−). Plants were then subjected to treatments with an excess of non-iron metals (-Fe/+++Metals; -/+++) as previously described (Dubeaux et al., 2018), or with Cd (-Fe/-Metals/+Cd; −/−/Cd): Plants were grown in climate-controlled growth chambers at 21°C under long day conditions (16h light/8h dark) with a light intensity of 120 μmol photon m^−2^s^−1^).

### Confocal imaging

For IRT1-mCitrine fluorescence, confocal imaging was performed using a Leica SP8 confocal laser scanning microscope. mCitrine was excited at 514nm and emission was collected from 530 to 580 nm. 5-day-old seedling grown in MS/2 without iron and non-iron metals medium (−/−) were treated as described in figure legends. For quantification, z-stacks encompassing the whole cell volume were generated. Stacks were then subjected to maximal projection and the mean fluorescence of whole cell and cytoplasm were quantified using ImageJ software with the tools “polygon selections”. Plasma membrane mean fluorescence were determined by subtraction of cell fluorescence and cytoplasm.

For meristems anatomy analyses and cell viability assays, 14-day-old seedlings were dark-incubated for 10min in a 10μg/ml Propidium iodide (PI) solution, rinsed and observed at the confocal microscope. PI was excited at 561nm and emission were collected from 590 to 650nm.

For FM4-64 endocytosis analyses, 5-day-old seedlings were dark-incubated in a 4μM FM4-64 solution for 10min, rinsed and observed after 30min to visualize endosomes. FM4-64 was excited at 488nm and emission were collected from 620 to 680nm.

### Root length and width measurement

Plants were grown vertically for 7 or 14 days, and root length and width determined using ImageJ software with the plugins “Segmentation” (tools “Simple Neurite Tracer”) and plugins “Straight”, respectively.

### Statistical analysis

All Statistical analyses were performed using GraphPad Prism 7.00 Software. Tests and details are indicated in figure legends.

### Western blot analyses

Total root proteins were extracted in Lammeli buffer using a 3:1 ratio (w/v). Protein detection was carried out using polyclonal anti-IRT1 (Agrisera, 1/5,000) and anti-rabbit horseradish peroxidase-coupled (BioRad, 1/20,000) antibodies. Detection of HRP chemiluminescence was performed with using SuperSignal West Dura Extended Duration Substrate (Thermo Scientific) on a ChemiDoc MP Imaging System (BioRad)

### Microscale thermophoresis

Binding experiments were performed by microscale thermophoresis with a Monolith NT.115 (NanoTemper Technologies, Munich, Germany). MBP fusions to the regulatory loop of IRT1 or IRT1_4HA_ were labelled with the Monolith NT™ Protein Labeling Kit RED according to the instructions provided by the manufacturer, using a 1:3 protein:dye molar ratio. For binding experiments, the labelled proteins (20nM) were incubated with a range of titrant concentrations (ZnCl_2_ or CdCl_2_) made by serial dilutions (1:2), in 50mM Tris buffer pH 7.4, 10mM MgCl_2_, 150mM NaCl, 0.05% Tween 20, in PCR tubes, at room temperature for 10 min. Premium treated capillaries (NanoTemper Technologies) were loaded and the measurements were performed at 25°C, 40% LED power and 40% microscale thermophoresis power, 20s laser-on time, 1s laser-off time. All the experiments were repeated at least twice with two independent protein labelling reactions. Binding data were analyzed using MO.AFFINITY ANALYSIS software (NanoTemper Technologies).

### Gene expression analyses

Total RNA was extracted from 50mg of root using the NucleoSpin^®^ RNA Kit (Macherey-Nagel). cDNAs were synthesized with the M-MLV Reverse Transcriptase (Promega) using 1μg of total RNAs and Oligo(dT). cDNA were 50-folds diluted and quantitative PCR were performed with a CFX Opus 384 Real-Time PCR System (BioRad). PCR amplification quality was visually controlled (melting and amplification curves) using the BioRad CFX Maestro software. Primers efficiencies were determined with the LinRegPCR software (Ruijter et al., 2009). Relative transcript level were normalized using At1g58050 and EF1a as described (Spielmann et al., 2020).

## Aknowledgements

We would like to acknowledge the Imaging facility from the Fédération de Recherche Agrobiosciences Interactions et Biodiversité of Toulouse (FRAIB) for access to microscopes and training. This work was supported by research grants from the French Laboratory of Excellence (project LabEx “TULIP” grant nos. ANR–10–LABX–41 and ANR–11–IDEX–0002–02) and the Fondation pour le Recherche Médicale (ECO20170637545).

---

**Figure S1:**
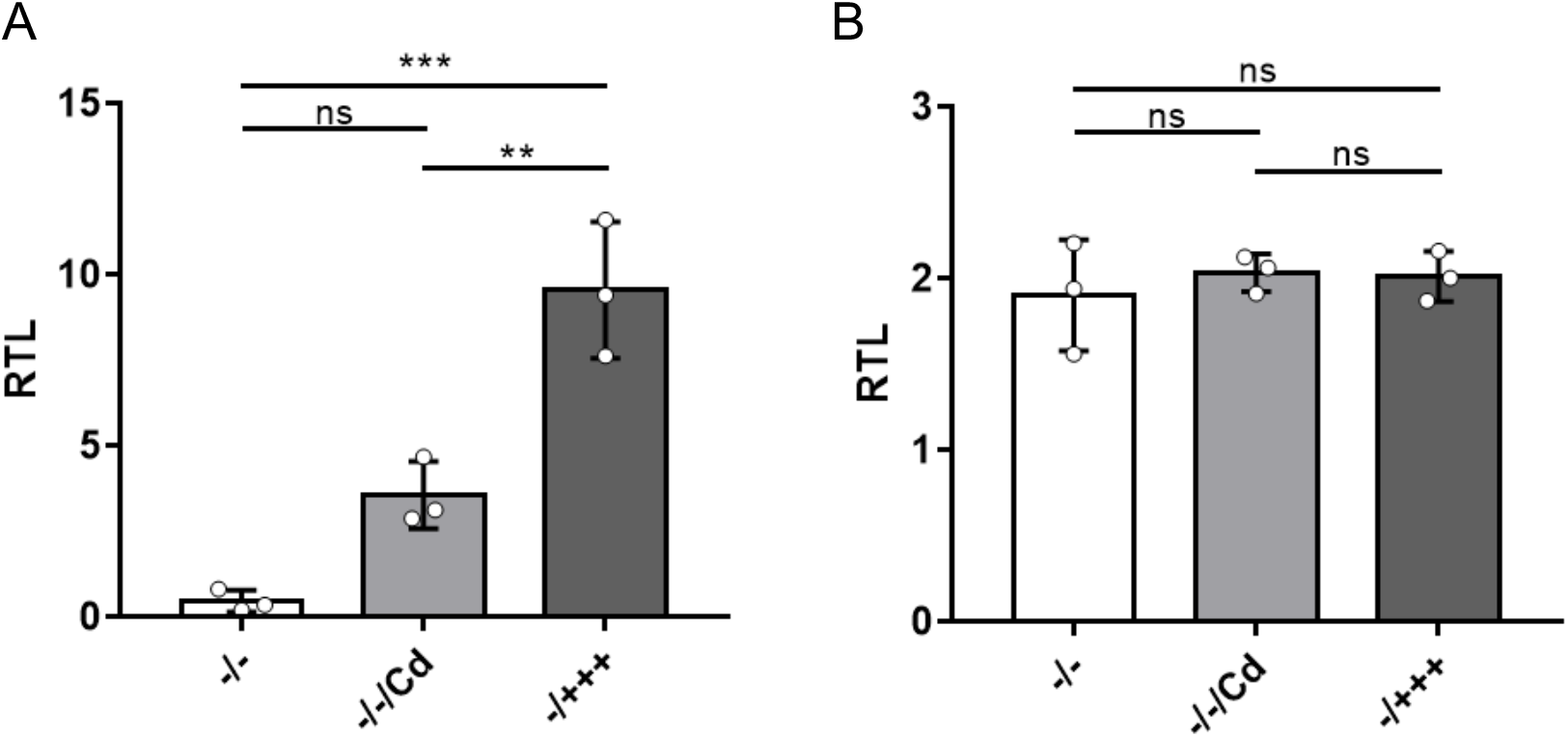
Influence of cadmium on *IRT1* transcription. Relative transcript level (RTL) of (A) IRT1 in Col-0 roots and (B) IRT1-mCit in roots of plants expressing PIN2::IRT1-mCit. Plants were grown in medium without iron and without non-iron metals (Zn, Mn and Co), then liquid- treated for 24h with the same medium (−/−), with cadmium 50μM (−/−/Cd) or with an excess of non-iron metals (−/+++). Value are means of 3 biological replicate and error bars represent SD. No significant differences between conditions were identify (one-way ANOVA, Tukey post-test; *** p < 0.001; ** p < 0.01; ns, not significant).

**Figure S2:**
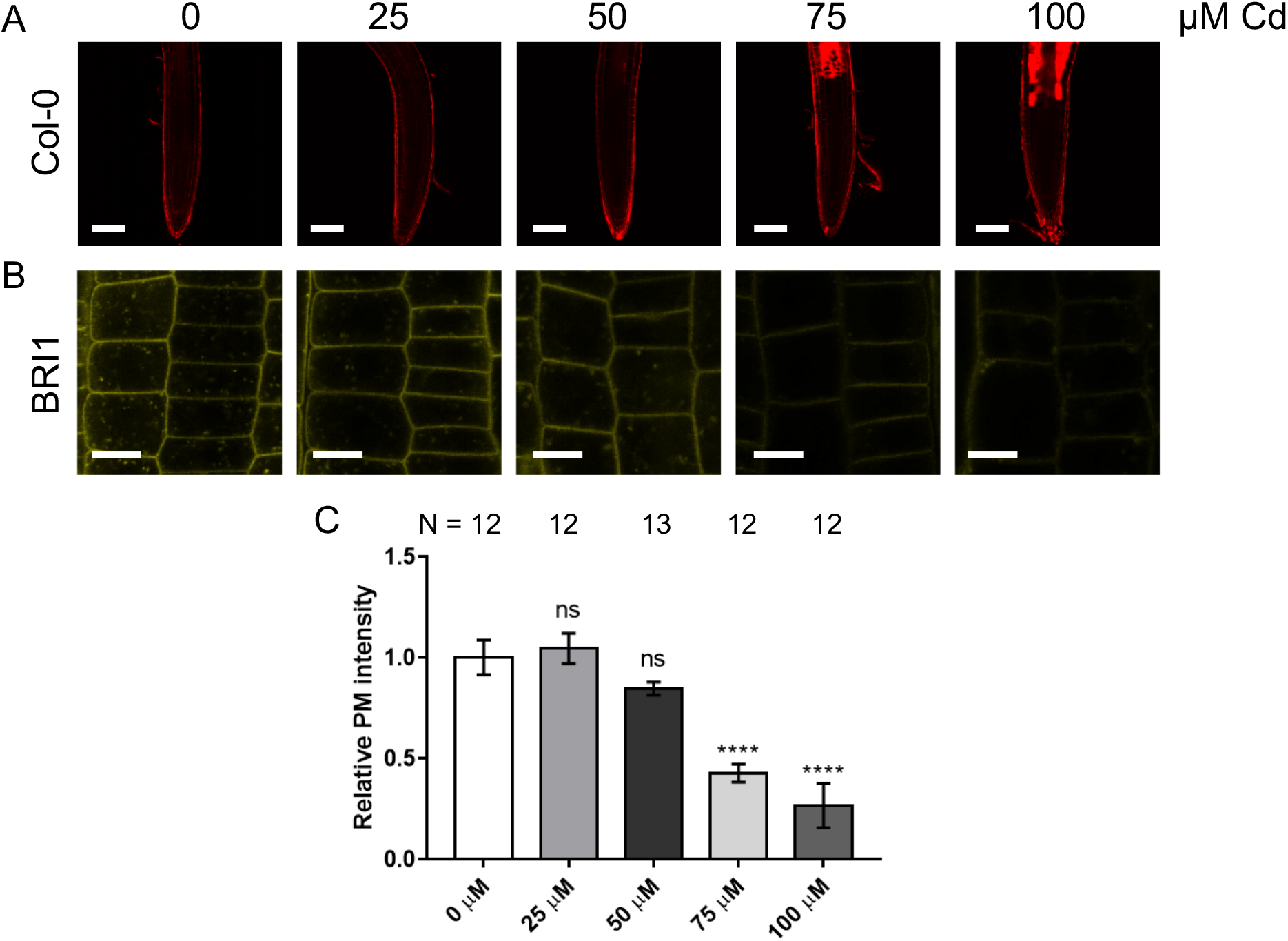
Assessment of cadmium toxicity. (A) Confocal microscopy analyses of Col-0 root tips treated with different cadmium concentrations. Propidium iodide (PI) was used to test cell viability. In living cells, PI is excluded from the cell and labels the cell wall. Dead cells show intracellular staining. Scale bars, 100μm (B) Confocal microscopy analyses of root epidermal cells from plants expressing BRI1::BRI1::mCit (BRI1) and exposed to different cadmium concentrations. Scale bars, 10μm (C) Quantification of the plasma membrane (PM) signal intensity of roots presented in (B). Value are relative to control condition (0μM Cd). Error bars represent SD and number of values are indicated in the graph. Asterisk indicate significant differences to control condition (0μM Cd) (One-way ANOVA, **** p < 0.0001; ns, not significant).

**Figure S3:**
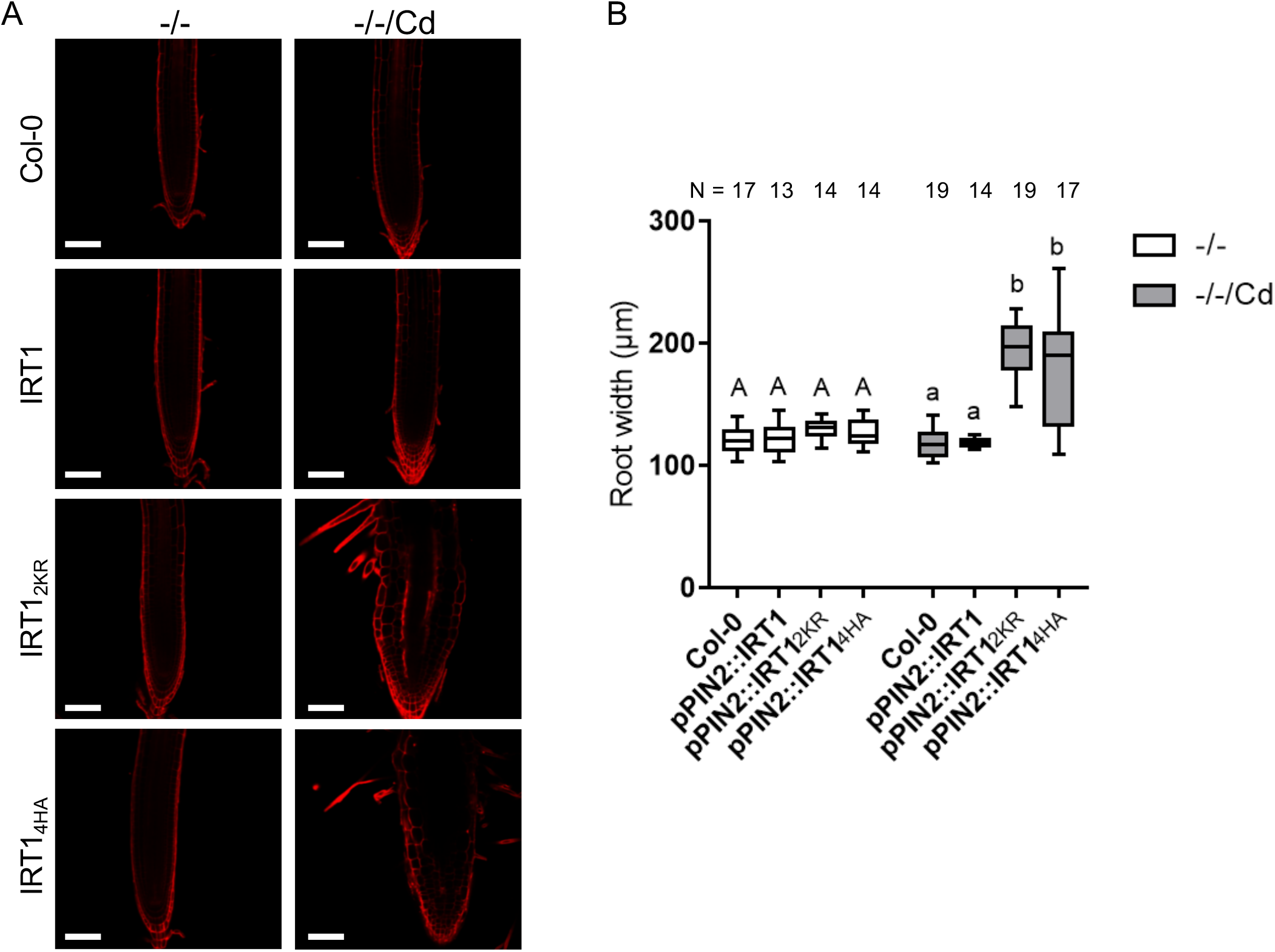
Analyses of meristem morphology. (A) Confocal microscopy analyses of root tips from Col-0 and plants expressing PIN2::IRT1-mCit (IRT1), PIN2::IRT1_4HA_-mCit (IRT1_4HA_) or PIN2::IRT1_2KR_-mCit (IRT1_2KR_) grown in absence of iron and non-iron metals (−/−) or in the same medium supplemented with 20μM of cadmium (−/−/Cd). Roots tips were stained with Propidium iodide. Scale bars, 100μm. (B) Determination of root width in the elongation zone of plant grown as in (A). Data are represented by Tukey boxplots and number of values are indicated in the graph. Different letters indicate significant differences between condition (Two-way ANOVA, Sidak post-test, p < 0.0001).

**Figure S4:**
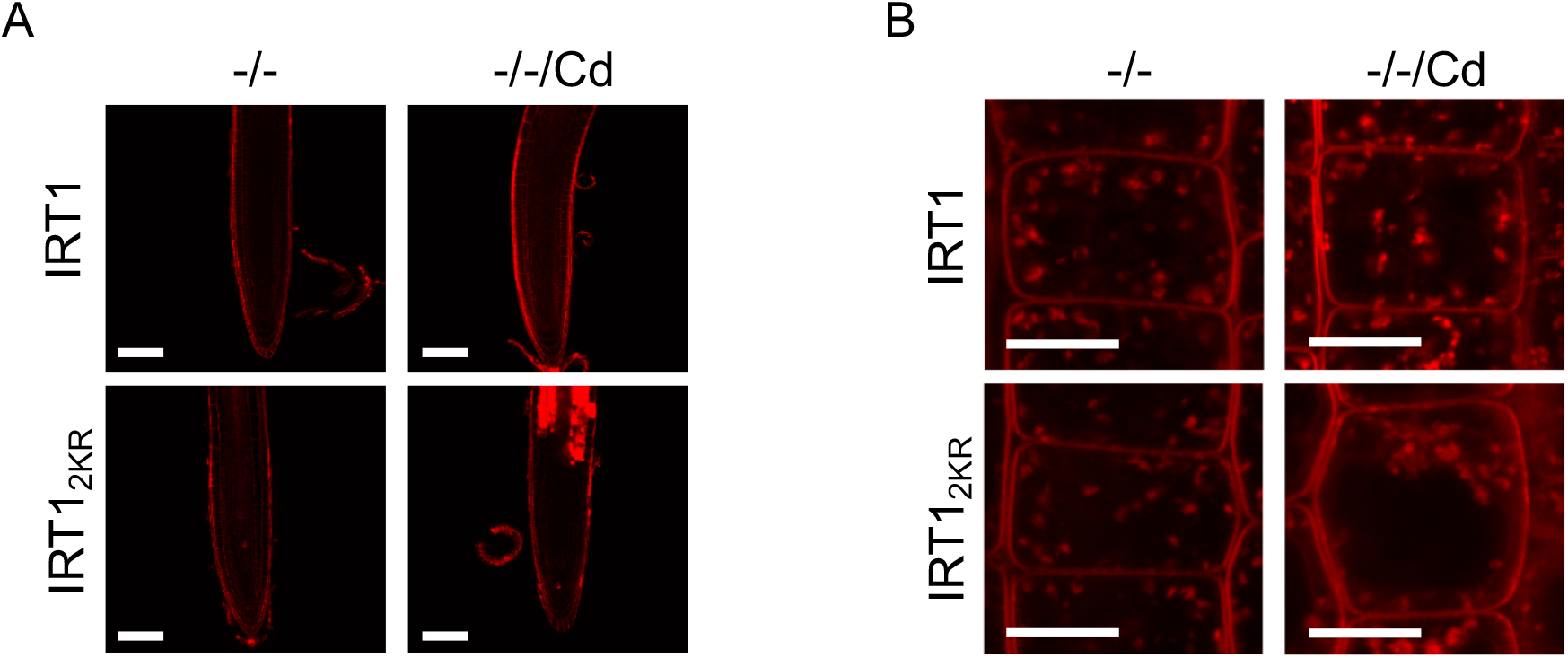
Influence of cadmium on cell viability and bulk endocytosis in IRT1_2KR_. Confocal microscopy analyses of root tips from plants expressing PIN2::IRT1-mCit (IRT1) or PIN2::IRT1_2KR_-mCit (IRT1_2KR_) and treated 24h with cadmium (−/−/Cd) or not (−/−). (A) PI was used to test cell viability. In living cells, PI is excluded from the cell and labels the cell wall. Dead cells show intracellular staining. Scale bars, 100μm. (B) FM4-64 staining of endocytosis in PIN2::IRT1-mCit (IRT1) or PIN2::IRT1_2KR_-mCit (IRT1_2KR_) treated 24h with cadmium (−/−/Cd) or not (−/−). Scale bars, 10μm.

